# A semi-automated tool for improving *Assessments* and *Lab Tests* data annotation in ImmPort

**DOI:** 10.64898/2026.10.02.756287

**Authors:** Amanda R. Tursi, Joshua D. Fortriede, Kevin Burns, Nicholas K. Saba, Parisa Pirani, Matthew D. Strub, Patrick Dunn, Henry Schaefer, Sandra Andorf

## Abstract

Public data sharing in immunological and clinical research is essential for transparency, reproducibility, and secondary analysis. However, many datasets in repositories such as ImmPort lack sufficient structured data to enable reuse. ImmPort’s *Assessments* and *Lab Tests* templates — which capture critical clinical information — are often incomplete or omitted due to their complexity and the manual effort required. To address this challenge, an application with a graphical user interface requiring no programming was developed to facilitate population of these templates. The tool streamlines the submission process by reducing time and complexity for data contributors while ensuring template formatting. It integrates ImmPort files, study-specific data files, and a curated data dictionary to automate the mapping of study variables to the required fields within each template. The tool was evaluated on seven ImmPort studies with diverse designs. It successfully generated submission-ready templates that passed ImmPort’s internal validation and will be incorporated into the repository. By improving the feasibility and completeness of templates, this approach enhances the quality, findability, and reusability of data within ImmPort.

## Introduction

Public data sharing within the biological sciences is largely recognized as crucial to promoting transparency, reproducibility, and enabling additional analysis of the data^1^. Despite these clear benefits, many researchers still fail to share their data or to reuse existing datasets^2,3^. Barriers for data depositors include a lack of knowledge about where and how to share their data, as well as reluctance to spend time cleaning and preparing datasets for public release^4–7^. Meanwhile, researchers seeking to utilize public data may be disincentivized by data incompleteness, poor annotation practices, and insufficient documentation^4,7^. Even if sufficient data are included in the repository, researchers may lack the computational skills to format and extract the relevant information they are seeking.

In particular, clinical trial data and the corresponding individual participant data (IPD) are often not shared in public repositories. When studies are shared, there are often quality issues, such as missing data and non-standardized formatting, that limit their reuse potential^8,9^. Compared to aggregate data, IPD enables more flexible and detailed meta-analyses, allowing researchers to explore different relationships and perform subgroup analyses that would otherwise be limited^10–12^. Secondary research based on IPD is considered the “gold standard” of systematic review, although successfully collecting this data for reuse remains a challenge^13–15^.

To address the numerous barriers to data sharing and reuse, high-quality, well-documented repositories are critical. For immunological and clinical data, a well-established open access platform is the Immunology Database and Analysis Portal (ImmPort), sponsored by the National Institutes of Health^16,17^. ImmPort contains over 1,300 clinical and mechanistic studies and is committed to following FAIR data principles for making data Findable, Accessible, Interoperable, and Reusable^18^. A key strength of ImmPort is its inclusion of IPD, case report forms (CRFs), data dictionaries, and protocols for many clinical studies.

Additionally, ImmPort promotes good documentation practices through guidelines and templates designed to standardize and annotate uploaded data, enabling data extraction for both single-study and multi-study analyses. The value of such structured data has been demonstrated in prior work by Chen et al., who used a uniformly curated dataset from multiple ImmPort studies to investigate post-donation outcomes in living kidney donors^19^. However, their curation process was performed manually, underscoring the labor-intensive nature of preparing data for reuse and highlighting the need for tools that can streamline large-scale data standardization^19^.

Indeed, complete annotation of uploaded datasets is often not fully achieved in ImmPort, largely due to the complexity of its template requirements. Instead, many individual data files are shared in an unstructured flat format, limiting their utility for secondary analyses. This is particularly pronounced in large, multifaceted clinical trials that incorporate diverse data types. Due to the time and effort required, data submitters often skip completing all relevant ImmPort templates, thereby failing to fully annotate their data for integration into the repository’s structured database. In particular, the *Assessments* and *Lab Tests* templates show low completion rates, with only 29.9% and 50.3% of clinical trials including this data, respectively (ImmPort data release 60; Figure 1). Notably, even when the *Assessments* and *Lab Tests* templates are submitted, this does not necessarily indicate that all relevant and available information has been incorporated in them.

**Figure 1.**
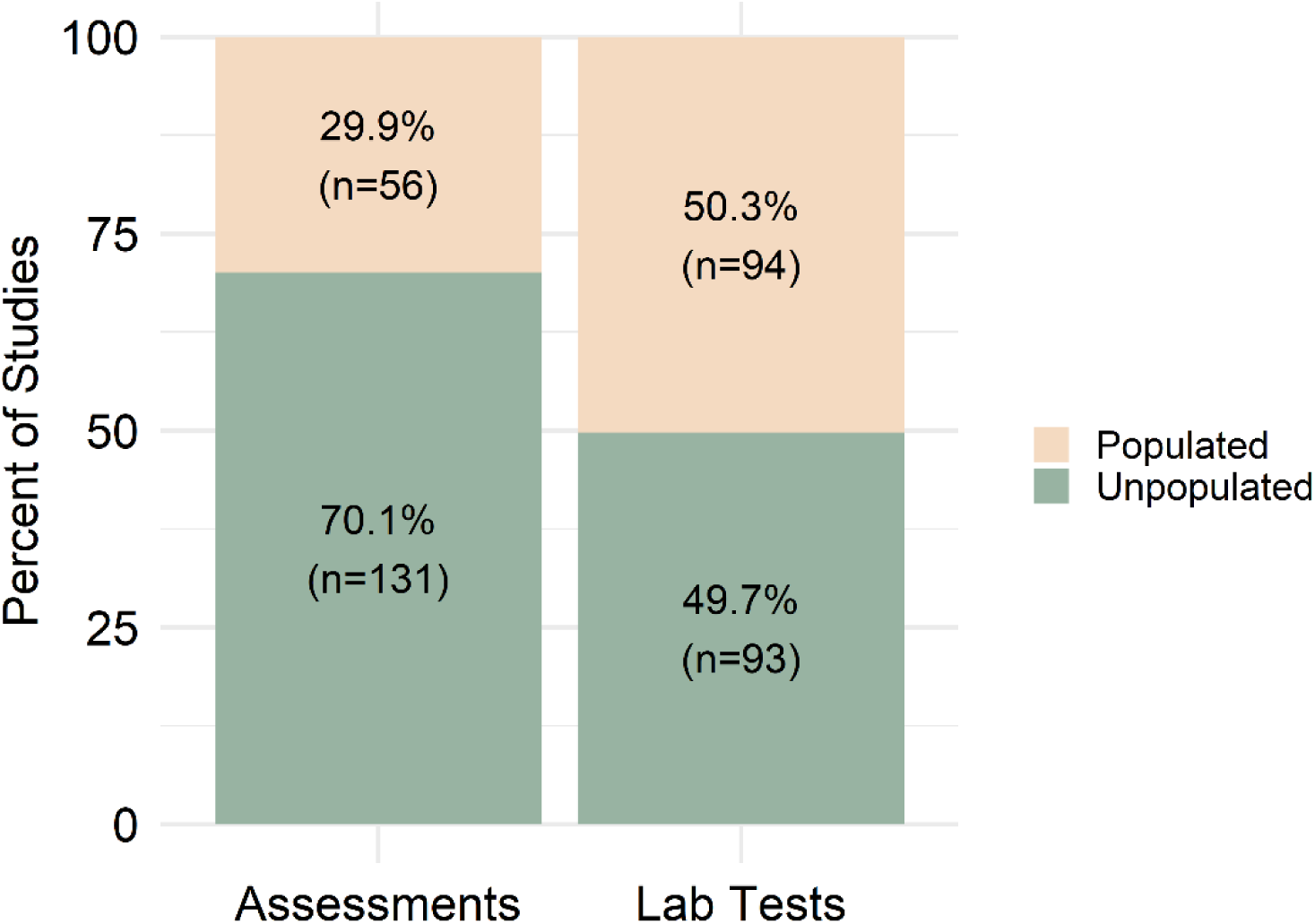
Percentage and number of ImmPort clinical trial studies that have at least one entry in the processed *Assessments* and *Lab Tests* tables (ImmPort data release 60).

Lab tests (e.g., complete blood count, metabolic panel, and urinalysis) and the non-sample-based results that encompass assessments (e.g., questionnaire responses, medical history, and skin prick test) contain vital information about the study subjects and outcomes. This information is often difficult to extract from the original study-specific data files, data dictionary, and CRFs, especially without programming expertise. Accordingly, ImmPort’s templates for these data types are complex. For example, a 20-question medical history survey generates an *Assessments* template with 327 columns, while a 14-test metabolic panel generates a *Lab Tests* template with 71 columns. Meanwhile, the number of rows scales with additional factors, such as the number of subjects, which encompass hundreds in many studies. As a result, the complexity and time-consuming nature of ImmPort’s data annotation process requires a solution to encourage both uploader compliance and researcher reuse.

The ImmPort Data Curation Tool presented here addresses these challenges by simplifying the annotation process through usage of a graphical user interface (GUI) that partially automates template production. This tool allows the user to upload their own study-specific files (data and results specific to their study), ImmPort files that were created in the earlier stages of the upload process, and a curated data dictionary (Figure 2A and 2B). The study-specific files and ImmPort files can be gathered with minimal manual effort, although the data dictionary requires additional curation. Nevertheless, the inclusion of a data dictionary is already expected in clinical trial submissions as part of good clinical data-sharing practices, which should limit the manual workload^20^. Once these files are loaded into the tool, only minimal input is required within the interface before submission. The tool then generates completed *Assessments* and/or *Lab Tests* templates, which the user can upload to ImmPort during the study creation and submission process (Figure 2C).

**Figure 2.**
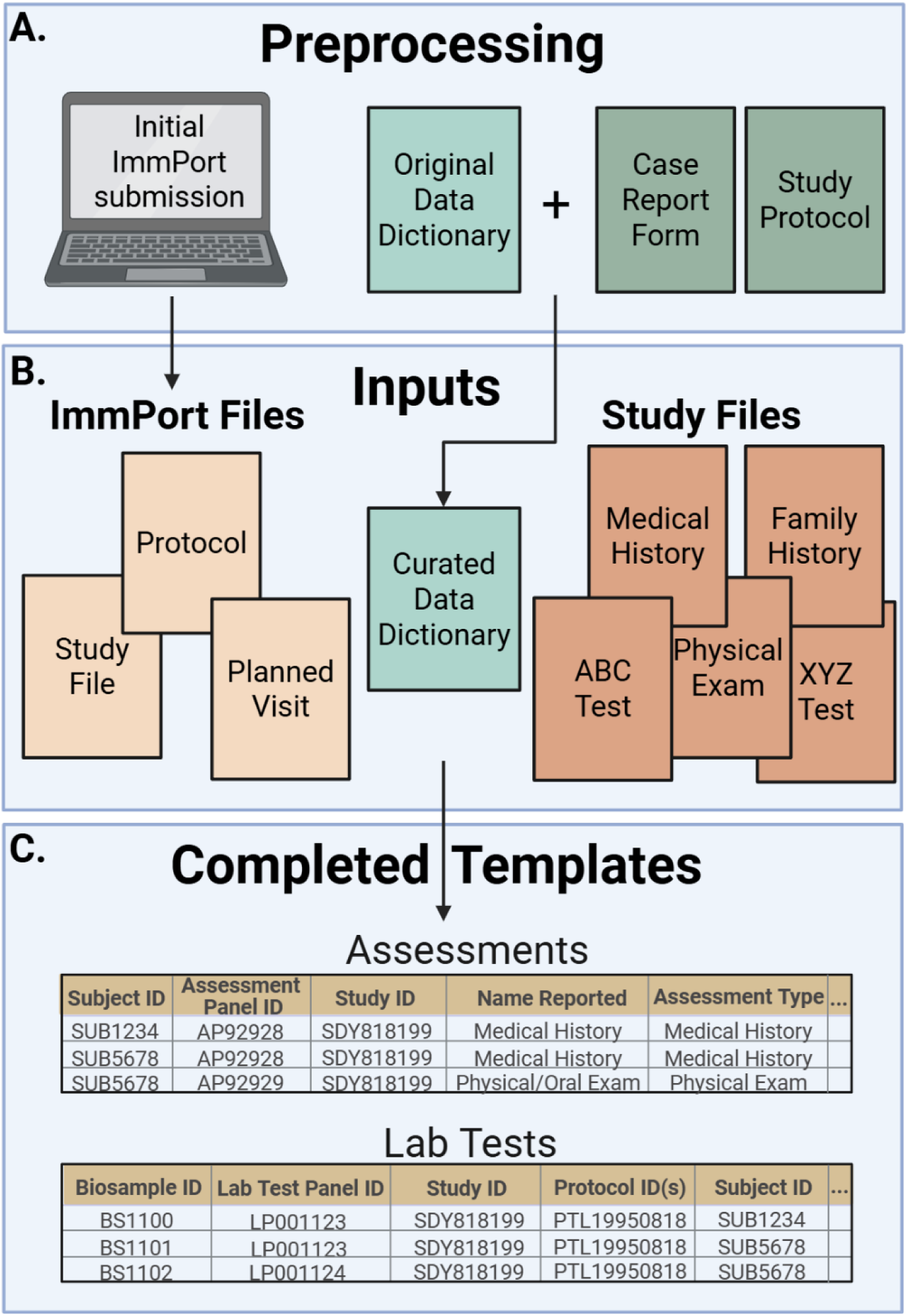
Overview of the tool’s file A) preprocessing requirements, B) inputs (ImmPort files, data dictionary, and study files) and C) template outputs (*Assessments* and/or *Lab Tests* templates).

By reducing the burden of template completion, the ImmPort Data Curation Tool facilitates more complete and consistent data annotation, ultimately enhancing the quality and reusability potential of studies deposited into ImmPort. In this work, the design and functionality of the tool is described, and its application is demonstrated with existing ImmPort datasets.

## Methods

### Tool Overview

The ImmPort Data Curation Tool was developed to partially automate completion of the *Assessments* and *Lab Tests* templates for data submission into the ImmPort repository. The tool requires preprocessing steps — most notably the curation of additional fields within an existing data dictionary — before it can be used. Once the data has been prepared, it can be uploaded into the tool, which contains 3 upload steps — “1. ImmPort Files”, “2. Data Dictionary”, and “3. Study Files”.

After these files are uploaded and the GUI fields are completed, the tool automatically generates and validates the filled templates, producing files that may contain thousands of rows and columns. These output files are ready for direct upload to ImmPort, where they are processed and made publicly available for reuse to the broader research community.

### Preprocessing Requirements

Before using the tool, several preprocessing steps must be completed to prepare the data for upload. These include study-specific data file collection, study registration and initial file upload, and data dictionary curation. The first two steps are standard requirements for ImmPort study submission, while the data dictionary curation is an additional requirement for the tool.

First, study-specific data files need to be gathered. Study files consist of any relevant IPD data collected from CRFs for that study. For example, one study file may contain liver function test results, while another may contain data from a neurological examination. In many clinical trials, multiple study files are generated from different CRFs through electronic data capture systems and are organized together within a single study folder.

Next, the user must begin the ImmPort study registration and upload process. This is necessary because, by ImmPort’s design, *Assessments* and *Lab Tests* templates must be filled in after broader templates, such as the protocol and study visits, have already been completed and uploaded. Uploading these initial templates and providing the study files are required steps for any ImmPort study submission, regardless of whether this tool is used. In other words, they require minimal additional effort to prepare for the ImmPort Data Curation Tool.

The final preprocessing step, curating the data dictionary, does require additional effort. While many aspects of the dictionary are optional for the user, the dictionary is essential, serving as the core component that links study file fields to ImmPort-compliant columns. As a result, completing the optional fields is encouraged when relevant, as doing so improves the quality of the generated templates.

### Data Dictionary Curation

While data dictionaries are routinely used in clinical trials, some curation is necessary for this tool (Figure 3). The additional curation provides important information, such as units or dates, that are incorporated into the final ImmPort templates. The first four columns of the dictionary — “Table Name”, “Field Name”, “Field Description”, and “Code List Values” — are often part of standard data dictionaries and are required for this tool. “Table Name” identifies the source study file, “Field Name” specifies the field within that file, “Field Description” provides a human-readable interpretation of that field, and “Code List Values” defines a mapping of coded responses to descriptive terms (e.g., 1 = Yes, 2 = No) for values within the field.

**Figure 3.**
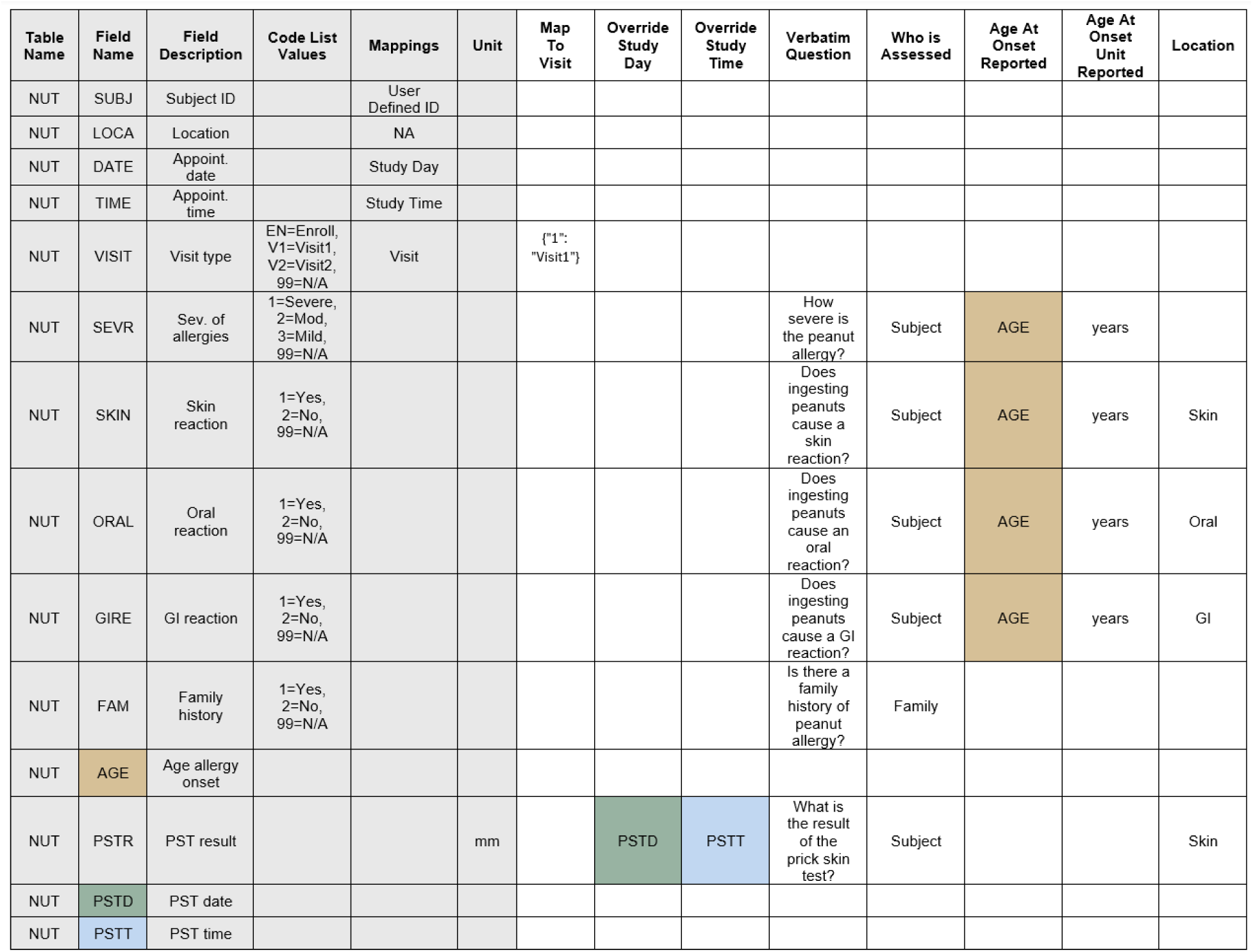
An example of a curated data dictionary populated with mock data to illustrate the required formatting. The first six columns (shaded in grey) represent required columns, while the remaining columns are optional but recommended when applicable. Green, blue, and tan colored entries indicate values that are mapped from entries listed in the “Field Name” column. For example, for the fields “SEVR”, “SKIN”, “ORAL”, and “GIRE”, “Age At Onset Reported” will be filled with values from the “AGE” field.

Two additional required columns –— “Mappings” and “Unit” — support integration with the ImmPort template schema. “Mappings” should contain categorical assignments that specify each field’s role in the study (e.g., “User Defined ID”, “Study Day”, “Study Time”, “Visit”, “Category”, or “NA”), whereas the “Unit” column can contain either a fixed unit (e.g., kg and percentage) or the name of another “Field Name” from which unit information is extracted. Additional information about how categorical assignments should be made in the “Mappings” column are available in the Supplement.

Additional optional columns — “Map To Visit” and “Override Study Day” — further enhance data mapping flexibility. “Map To Visit” can contain visit identification information that is encompassed in the corresponding study file field but is not congruently named in the planned_visit ImmPort file. The “Override Study Day” column should be used when different fields within a study file are associated with different days, making the global mapping “Study Day” in the “Mappings” column insufficient. In these cases, users can assign a global mapping in the “Mappings” column and provide more granular overrides in “Override Study Day”. Overrides may reference days (in integer format) or another “Field Name” that points to a study file field that contains the override day information. For example, in Figure 3 the field “DATE” is assigned as “Study Day” in the “Mappings” column. If there were no values filled in the “Override Study Day” column, all fields would have the “DATE” day assigned. However, in this example, the “PSTR” field has “Override Study Day” filled in as “PSTD”, another field in the study file (shown in green). As a result, “PSTR” will have the values from “PSTD” assigned and all other fields will have the values from the “DATE” field denoted.

The final columns —– “Override Study Time”, “Verbatim Question”, “Who is Assessed”, “Age at Onset Reported”, “Age at Onset Unit Reported”, and “Location” — are also optional and only relevant to the *Assessments* Template. The “Override Study Time” column works similarly to “Override Study Day”, only it overrides times instead of days. The “Verbatim Question” should indicate the question specified on the CRF, while “Who is Assessed” should indicate who each field refers to. “Age at Onset Reported” and “Age at Onset Unit Reported” refers to the age of onset and the corresponding unit for that field, respectively. Finally, “Location” should indicate the organ or body system the field indicates. As is the case with the “Unit” and “Override Study Day” columns, these columns can either be filled manually or reference another “Field Name” in the data dictionary.

### Data Processing Workflow

The tool integrates information from four primary sources: (1) Files generated during earlier stages of the ImmPort submission process, (2) a curated data dictionary, (3) study-specific data files, and (4) information provided by the user within the GUI (Tables S1 and S2). The two ImmPort templates — *Assessments* and *Lab Tests* — are mutually exclusive and structured in a wide format. Each template includes a set of static columns followed by repeating blocks. Specifically, the *Assessments* template consists of seven static columns and sixteen recurring columns, while the *Lab Tests* template includes fifteen static columns and four recurring columns. Within both templates, certain columns must be fully populated, in compliance with ImmPort’s requirements, while others are optional and can be left blank if irrelevant or if the information is not available. Depending on the number of fields in the study-specific data files, the completed templates can easily contain hundreds — or even thousands — of populated columns per study file.

To reduce manual effort, the tool semi-automates the mapping of data into template columns. First, users upload ImmPort-generated files — study_file.txt, protocol.txt, and planned_visit.txt — which are produced after the initial study registration (Figure 4). These individual files can be downloaded from ImmPort, either before or after they have been published on the site. Alternatively, a ZIP file (called the “Tab” file in ImmPort) containing the three files can be downloaded after publication. These templates are thenparsed by the tool to aid in populating template columns such as “Study ID”, “Planned Visit ID”, “Protocol ID(s)” (in *Lab Tests*), and “Name Reported” (in *Assessments*).

**Figure 4.**
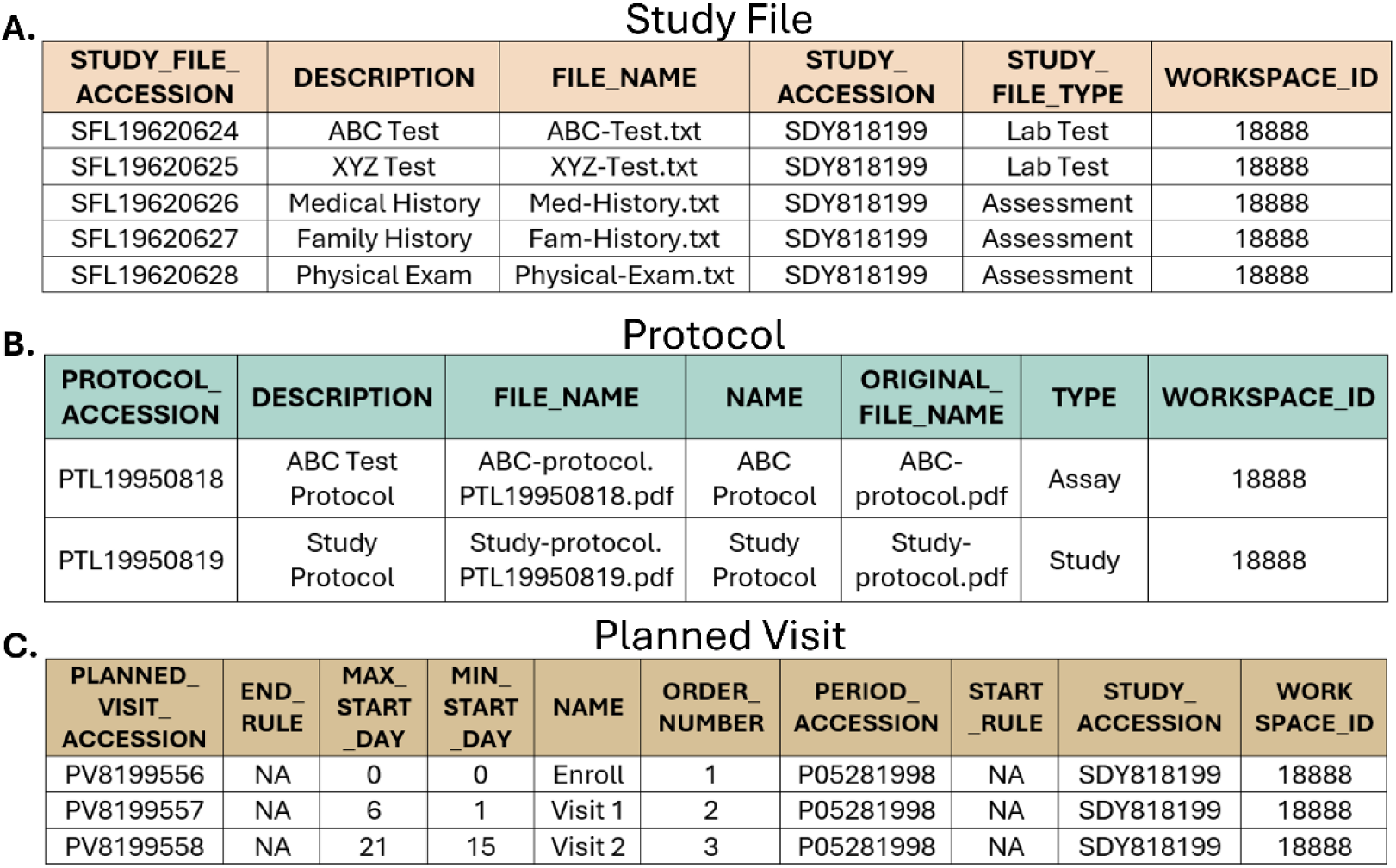
The required ImmPort-generated input files for the data curation tool, shown in the post-publication format. The A) study file, B) protocol, and C) planned visit tables are displayed with the expected column structure and populated with mock data.

Next, the curated data dictionary is used to interpret fields within the study-specific data files to aid in populating additional template columns, such as “Subject ID”, “Result Value Reported”, and “Result Unit Reported”. The dictionary also helps fill in template-specific columns, including “Name Reported” (in *Lab Tests*) and “Age At Onset Reported”, “Age At Onset Unit Reported”, “Location Of Finding Reported”, “Verbatim Question”, and “Who Is Assessed” (in *Assessments*). This dictionary provides human-and machine-readable mappings, enabling standardized interpretation of fields within the study files, as well as additional information that can be used to fill out the templates.

After the ImmPort-generated files and the data dictionary are uploaded into the tool, the study-specific data files themselves can be uploaded. This tool supports uploading an entire folder of individual study files for further processing with subsequent user input via the GUI to support template generation. For each uploaded study file, users may define a default planned visit if one is not specified in the data. The GUI also allows manual or dropdown selection of relevant template columns such as “Assessment Name” (for *Assessments*) and “Protocol”, “Type”, “Subtype”, “Name Reported”, “Study Time T0 Event”, and “Study Time T0 Event Specify” (for *Lab Tests*). The “Protocol” options are populated from the uploaded protocol.txt file, while the “Type”, “Name Reported”, and “Study Time T0 Event” options are derived from preexisting ImmPort options. When none of the available choices for “Type” or “Study Time T0 Event” accurately describe the data, users may select “Other” and provide a custom “Subtype” or “Study Time T0 Event Specify”, respectively.

After all other columns in the templates have been populated, the curation tool assigns unique accession numbers that will later be converted by ImmPort into ImmPort-specific accessions. For the *Lab Tests* template, “Biosample ID” is assigned for each unique combination of “Subject ID”, “Planned Visit ID”, and “Type”, while “Lab Test Panel ID” is assigned a unique value for each study file. Likewise, “Assessment Panel ID” is assigned a unique value for each study file in the *Assessment*s template.

Throughout the uploading and processing workflow, the tool performs validation checks to ensure compliance with tool and ImmPort requirements. When there is missing mandatory data, invalid mappings, or formatting inconsistencies, the tool displays descriptive error messages to prompt the user to correct issues for accurate template generation.

### Tool Implementation

The ImmPort Data Curation Tool was written in Python (version 3.13.1) and run in a Jupyter Notebook environment (version 6.2.0)^21^. A GUI was implemented using the ipywidgets (version 7.8.5) and ipyfilechooser (version 0.6.0) libraries to facilitate user interaction^22,23^. It was rendered as a standalone web application using Voilà (version 0.5.0)^24^. Core data processing and manipulation were performed using pandas (version 2.2.3) and NumPy (version 2.2.1)^25–27^.

### Use Case Demonstration

To evaluate tool performance and demonstrate its applicability across diverse datasets, we applied it to seven studies from the ImmPort repository (Table 1). For each study, the ImmPort Tab ZIP file and StudyFiles folder were downloaded from ImmPort, along with associated CRFs, protocol forms, and study descriptions (ImmPort data release 60). Study files were manually reviewed and assigned to either the *Assessments* or *Lab Tests* template based on purpose and content. Files deemed extraneous — such as those containing administrative data or information better suited to other ImmPort templates (e.g., adverseEvents.txt, subjectHumans.txt, and interventions.txt) — were excluded.

**Table 1.** ImmPort studies used to demonstrate the curation tool, including the study accession number, study name, and the number of study files transformed into the *Assessments* and *Lab Tests* templates. The total number of study files and average (± standard deviation) across the studies per template are also reported.

| ImmPort Accession | ImmPort Study Name | # of Study Files for Assessments Template | # of Study Files for Lab Tests Template |
| --- | --- | --- | --- |
| SDY620<sup>28</sup> | Atopic Dermatitis Research Network (ADRN) Influenza Vaccine Study (ADRN-05) | 8 | 6 |
| SDY1027<sup>29</sup> | The Role of Epigenetics in Inner City Asthma (ICAC-15) | 12 | 5 |
| SDY1433<sup>30</sup> | Optimization of NULOJIX (Belatacept) Usage as a Means of Minimizing CNI Exposure in Simultaneous Pancreas and Kidney Transplantation (CTOT-15) | 13 | 9 |
| SDY1515<sup>31,32</sup> | Peanut Sublingual Immunotherapy (CoFAR4) | 14 | 4 |
| SDY1519<sup>33,34</sup> | Eosinophilic Esophagitis Databank (CoFAR5) | 11 | 2 |
| SDY1520<sup>35,36</sup> | Peanut Epicutaneous Immunotherapy (CoFAR6) | 14 | 5 |
| SDY1550<sup>37</sup> | Baked Egg or Egg Oral Immunotherapy for Children With Egg Allergy (CoFAR7) | 15 | 5 |
| **Total** |  | 87 | 36 |
| **Average (± Standard Deviation)** | | 12.43 (± 2.37) | 5.14 (± 2.12) |

Each study’s existing data dictionary was reformatted to comply with tool requirements and additional columns were populated with information from ImmPort (e.g., CRFs and protocol forms). The curated dictionary, Tab ZIP file, and study files were then uploaded to the tool. Appropriate options were selected in the GUI, and the resulting *Assessments* and *Lab Tests* templates were generated. The outputted files were manually reviewed to ensure alignment with ImmPort formatting standards. Following this review, they were submitted to ImmPort, where they underwent internal validation to confirm compliance with repository requirements. All finalized templates will be publicly shared by ImmPort in a future data release.

### Example Dataset Creation

To provide an illustrative example, an anonymized dataset was created that was derived from ImmPort study SDY1550 (CoFAR 7: baked egg or egg oral immunotherapy trial)^37^. Six study files were selected to demonstrate various features in the curated data dictionary and the tool’s ability to propagate both templates. Four files (*Medical History 1*, *Medical History 2*, *Oral Food Challenge*, and *Skin Prick Test*) were used for the *Assessments* template, while two files (*Basophil Activation* and *Immunoglobulin E*) were used for the *Lab Tests* template.

To anonymize the data, all identifiers were randomized, categorical data were shuffled, and numerical values were altered within ±5% using random noise. Numerous fields were removed for simplicity. The final example set includes a curated data dictionary and mock ImmPort files (study_file.txt, protocol.txt, and planned_visit.txt) that match the modified study files. This example, along with a step-by-step tutorial, is distributed with the tool as a reference for users.

### Ethics Statement

This study did not involve the collection of new human data. All data used were obtained from ImmPort and derived from previously published studies that had received ethical approval from their respective institutional review boards.

## Results

### Tool Interface Overview

The GUI was designed to streamline the data upload and annotation process. It includes three data upload sections. The first is the “1. ImmPort Files” section, where the three previously produced ImmPort files are uploaded (Figure 5). These files contain vital information including the study accession, the planned visits, and protocol information. The files can be uploaded individually (Figure 5A) or combined in a ZIP file (Figure 5B). If an uploaded file does not abide by the ImmPort required format (Figure 4), an error message will appear telling the user to ensure their file is compliant before proceeding. Additionally, the user can optionally specify the study accession in the study ID textbox if the uploaded study_file does not include that information.

**Figure 5.**
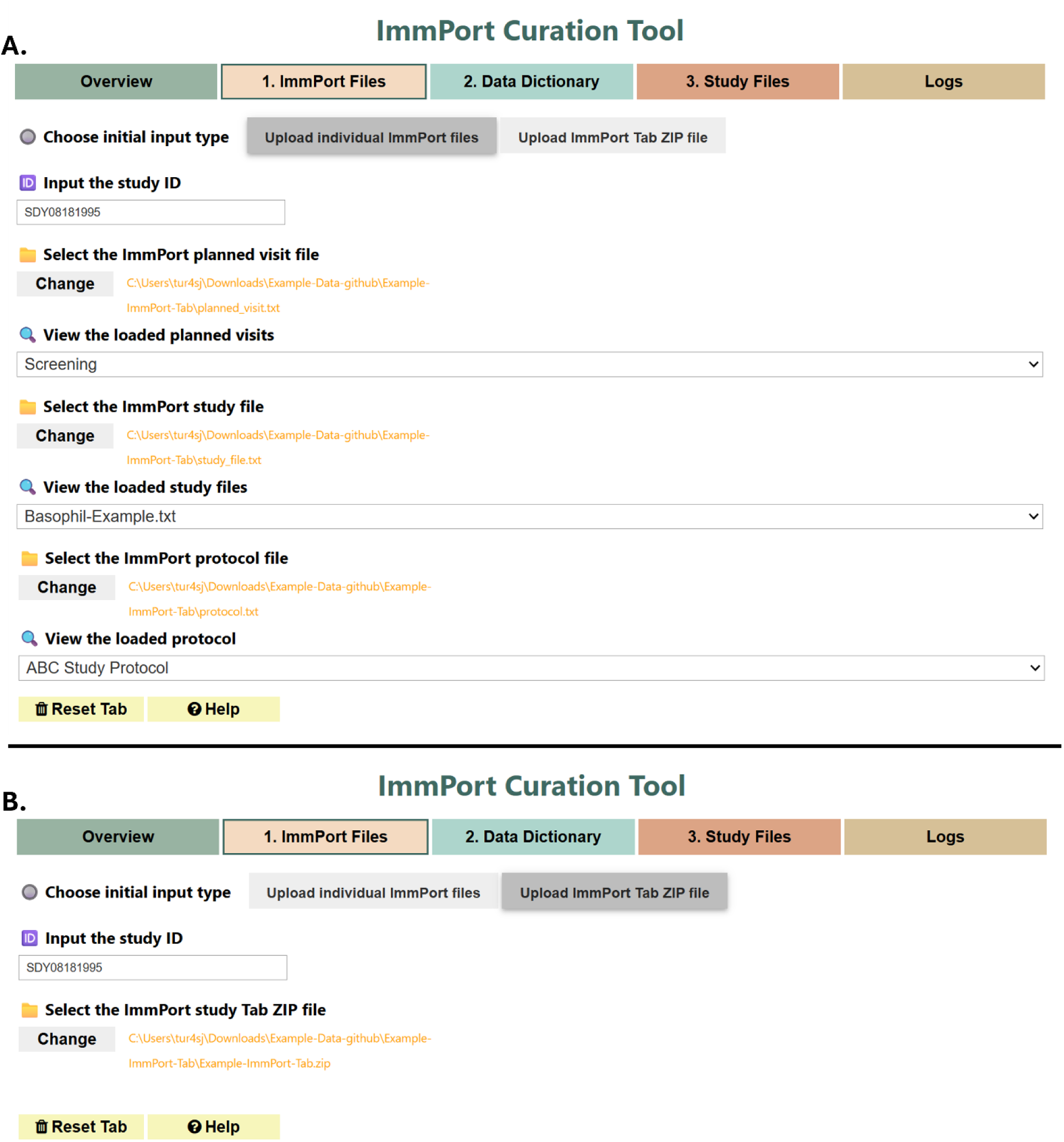
Screenshot of the “1. ImmPort Files” tab in the GUI of the curation tool. This tab allows users to upload the previously created ImmPort files. A) the three required files (planned_visit, study_file, and protocol) can be uploaded individually or B) they can be uploaded together in a ZIP file. Example data is shown in the application for demonstration purposes.

The second upload section is the “2. Data Dictionary” section, where the curated data dictionary is uploaded (Figure 6). The data dictionary is vital to mapping the study data to template columns. This section also allows the user to specify whether they plan to generate *Assessments* templates, *Lab Tests* templates, or both. Again, the tool contains an internal validation to ensure the required data dictionary format has been met. If not, the tool will display an error message indicating that there is an issue with the uploaded file.

**Figure 6.**
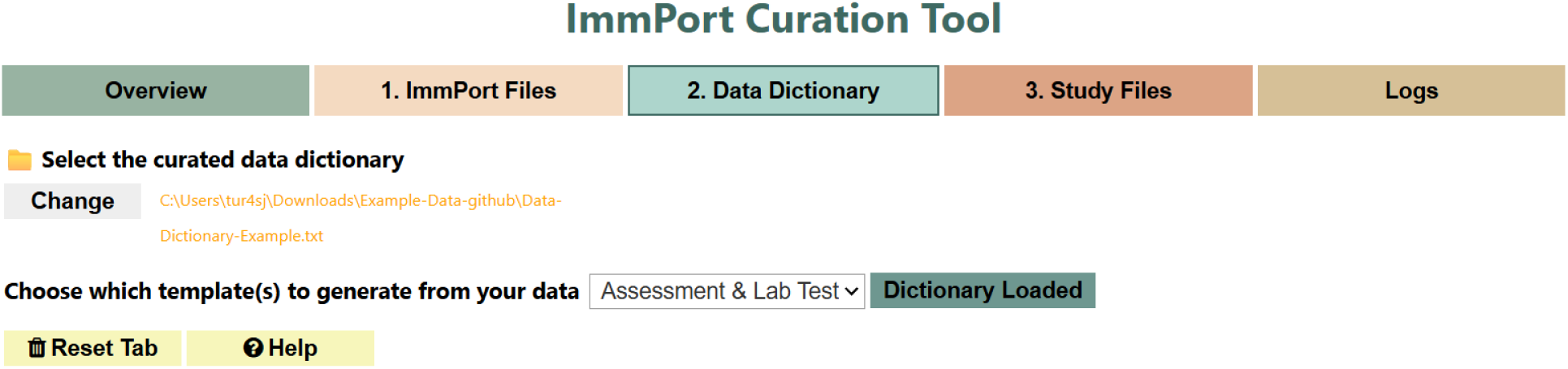
Screenshot of the “2. Data Dictionary” tab in the GUI of the curation tool. This tab allows users to upload the curated data dictionary and specify the template type(s) they plan to create. Example data is shown in the application for demonstration purposes.

The third and final upload section is “3. Study Files”, where the study-specific data files are provided (Figure 7). In this section, a directory containing the study files must be selected, and these files should correspond to information in the previously uploaded files from the first two sections. Additional details for each study file can be entered within the interface. Once this step is complete, templates can be generated. All uploaded files and entered information are combined and parsed to create the final templates. If an error occurs during template generation, the user will be notified.

**Figure 7.**
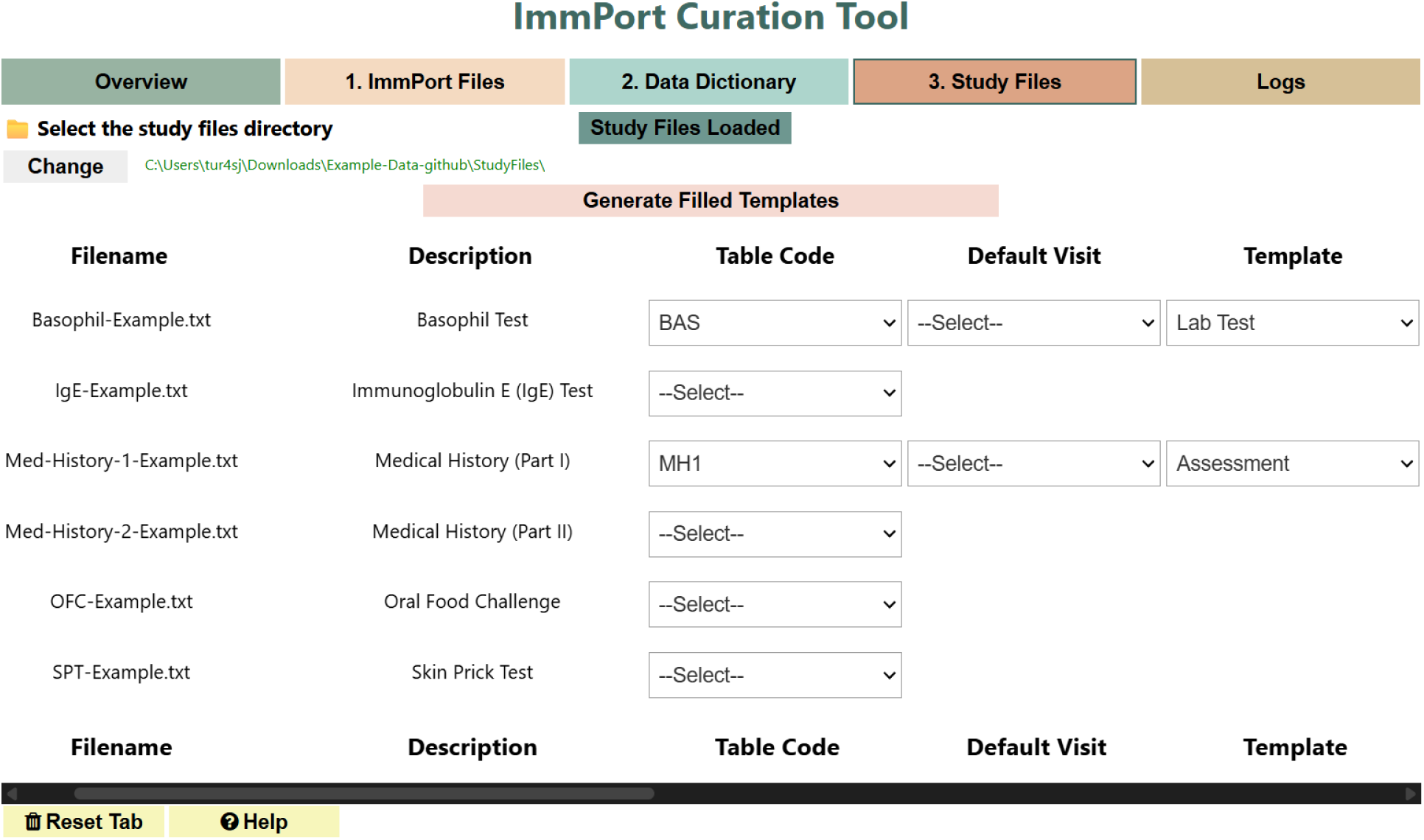
Screenshot of the “3. Study Files” tab in the GUI of the curation tool. This tab allows users to upload the study-specific data files that will be transformed into the *Assessments* and/or *Lab Tests* templates. Example data is shown in the application for demonstration purposes. Note that only a portion of the available user options are shown in this view.

Additionally, the tool contains the more general “Overview” section that provides a summary of the tool’s functionality, along with links to more expansive documentation (Figure S1). This section also includes an example data download link, which allows users to try the tool alongside a step-by-step tutorial. Finally, the tool contains a “Log” section, which displays real-time status updates and error messages that can assist users with troubleshooting the application (Figure S2). All alterable sections also include help and reset buttons to enhance usability.

### Evaluation Across Seven ImmPort Studies

To evaluate the performance and versatility of the ImmPort Data Curation Tool, it was applied to seven publicly available studies from the ImmPort repository. These studies were selected to represent a diverse range of research consortia, study designs, and data structures. Specifically, one study (SDY1433) was part of the Clinical Trials in Organ Transplantation (CTOT) program, one study (SDY620) was from the Atopic Dermatitis Research Network (ADRN), one study (SDY1027) was part of the Inner City Asthma Consortium (ICAC), and four studies (SDY1515, SDY1519, SDY1520, and SDY1550) were from the Consortium for Food Allergy Research (CoFAR)^28–37^. These studies contained between eight and fifteen (average: 12.43) study files that were suitable for the *Assessments* template and between two and nine (average: 5.14) study files that were suitable for the *Lab Tests* template (Table 1). In total, the studies encompassed 123 study files suited for the *Assessments* and *Lab Tests* templates, representing just over half (51.3%) of all study files uploaded to ImmPort across the seven evaluated studies (Table S3).

For each study, the tool was used to generate wide-format tables for both the *Assessments* and *Lab Tests* templates (Tables S4-S10). Each study file populated its own sub-template, which could then be uploaded to ImmPort where all *Assessments* or *Lab Tests* templates were combined into one final *Assessments* or *Lab Tests* template, respectively.

Across the seven studies, the final *Assessments* templates contained an average of 2,038.4 rows and 20,424.4 columns, while the final *Lab Tests* templates contained an average of 1,899.6 rows and 4,240 columns (Table 2). The relatively large standard deviations reflected substantial diversity in study size, study design, and the number and complexity of study files across the evaluated studies (Table 2). Specifically, the *Assessments* templates ranged from 474 to 6,295 rows and from 1,256 to 42,158 columns, whereas the *Lab Tests* templates ranged from 708 to 2,909 rows and from 158 to 16,637 columns (Table 2). Since some columns always remained empty in all rows, the number of filled columns was always fewer than the total number of columns. As a result, the median number of columns filled across all rows was also calculated to better reflect actual data density (Tables S4-S10). The median often remained in the hundreds or even thousands, underscoring the substantial amount of information contained in the templates.

**Table 2.** The total number of rows and columns generated for each ImmPort study after transformation into the final *Assessments* and *Lab Tests* templates. For each study, the final templates encompass all sub-templates combined. Across the studies per template, the total summation of rows and columns and average (± standard deviation) are also reported.

| ImmPort Study Accession | ImmPort Study Title | Assessments Template |  | Lab Tests Template |  |
| --- | --- | --- | --- | --- | --- |
|  |  | # of rows | # of columns | # of rows | # of columns |
| SDY620 | Atopic Dermatitis Research Network (ADRN) Influenza Vaccine Study (ADRN-05) | 1,808 | 1,256 | 1,635 | 298 |
| SDY1027 | The Role of Epigenetics in Inner City Asthma (ICAC-15) | 1,615 | 11,628 | 708 | 313 |
| SDY1433 | Optimization of NULOJIX (Belatacept) Usage as a Means of Minimizing CNI Exposure in Simultaneous Pancreas and Kidney Transplantation (CTOT-15) | 474 | 16,709 | 2,540 | 369 |
| SDY1515 | Peanut Sublingual Immunotherapy (CoFAR4) | 747 | 42,158 | 965 | 5,472 |
| SDY1519 | Eosinophilic Esophagitis Databank (CoFAR5) | 6,295 | 4,399 | 1,746 | 158 |
| SDY1520 | Peanut Epicutaneous Immunotherapy (CoFAR6) | 1,143 | 34,334 | 2,794 | 6,433 |
| SDY1550 | Baked Egg or Egg Oral<br>Immunotherapy for Children<br>With Egg Allergy (CoFAR7) | 2,187 | 32,487 | 2,909 | 16,637 |
| **Total** |  | 14,269 | 142,971 | 13,297 | 29,680 |
| **Average (± Standard Deviation)** | | 2,038.4<br>(± 1,970.2) | 20,424.4<br>(± 15,951.8) | 1,899.6<br>(± 877.1) | 4,240<br>(± 6,091.3) |

The dimensions of the propagated templates differed across studies and were largely the result of the individual study files containing differing numbers of patients, study visits, and questions in the CRFs. Both the completed *Assessments* and *Lab Tests* templates contain static columns — which only appear once — and recurring columns, which can repeat multiple times. As a result, the number of rows in the generated templates represented the number of unique entries in the static columns. Here, there were no clear patterns in the

*Assessments* verses *Lab Tests* templates and the variability across studies was more limited (Figure 8A). In contrast, the number of columns in the generated templates were largely reflective of the number of recurring columns and how frequently those recurring columns were repeated. The number of columns showed more variability across studies (Figure 8B). Notably, for all seven studies, the generated *Assessments* templates had more columns than the *Lab Tests* templates. This partially reflects the difference in the number of recurring columns between templates (sixteen columns for *Assessments* and four for *Lab Tests*).

**Figure 8.**
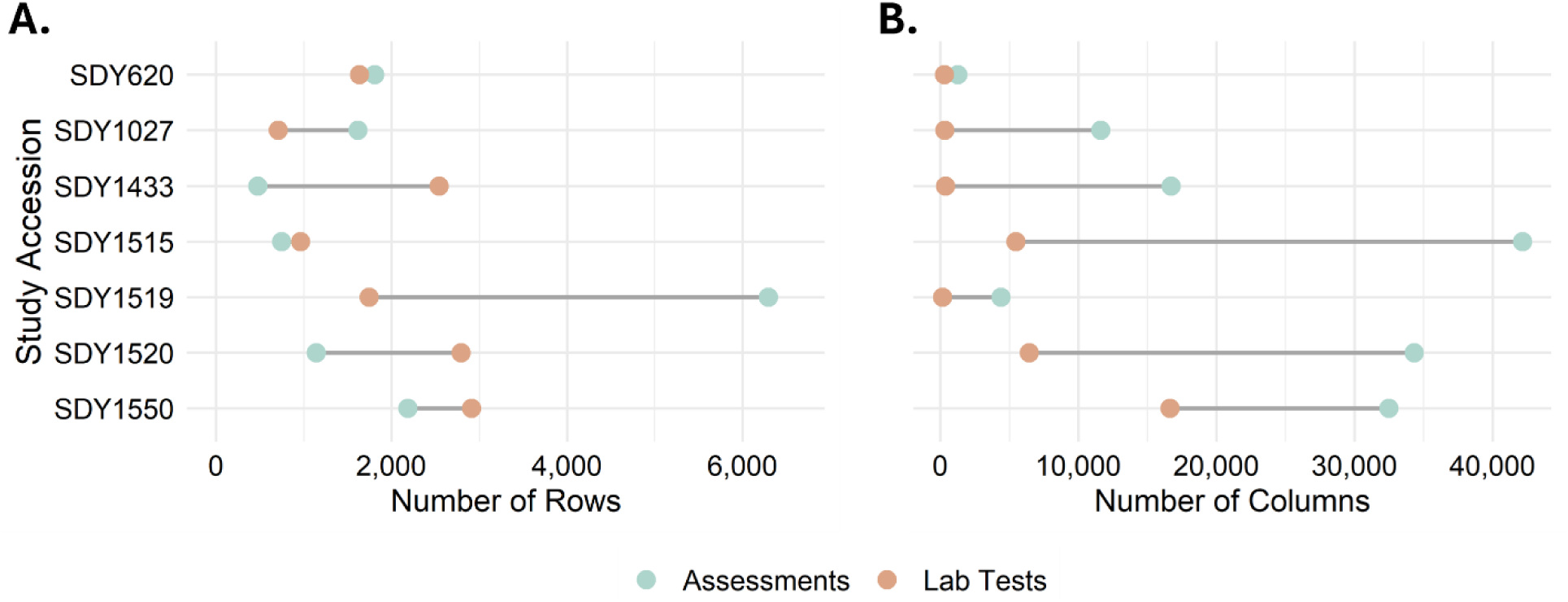
Dumbbell plots showing the evaluated ImmPort study accessions and the number of produced A) rows and B) columns across the final *Assessments* (blue) and *Lab Tests* (orange) templates. For each study, the final templates encompass all sub-templates combined.

The generated templates were manually reviewed to ensure consistency with ImmPort formatting standards. Following this review, the files were submitted to ImmPort, where they passed internal quality control checks and were accepted for integration into the repository. They will be incorporated into the repository’s structured database and made publicly available in a future data release. Specifically, the data will be transformed into standardized ImmPort files, called assessment_component, assessment_panel, lab_test, and lab_test_panel. These outputs will confirm that the tool successfully generated submission-ready templates that met ImmPort’s formatting requirements.

### Usability Evaluation

The ImmPort Data Curation Tool and the resulting templates were evaluated both by the ImmPort Curation Team and by researchers not associated with ImmPort. Testing was conducted across multiple operating systems to ensure compatibility.

Overall, participants reported that the tool was intuitive to use, with a well-organized GUI that guided users through the required workflow of uploading the ImmPort files, curated data dictionary, and study-specific data files before template generation. The built-in validation checks, help features, and logging provided useful guidance throughout the curation process and made it straightforward to identify and resolve input issues.

From a curation perspective, the tool demonstrated strong performance in generating templates that adhered to the required ImmPort structure. The generated *Assessments* and *Lab Tests* templates consistently maintained correct relationships between study metadata, subjects, visits, biosamples, and protocols while accurately preserving result values, units, and source terminology. The outputs required only minimal curator review before successfully passing ImmPort internal validation and repository quality control.

Participants noted that the preparation of the well-curated data dictionary was the most time-intensive step in the process. This was expected, as the quality and completeness of the generated templates depend on the accuracy of the mappings provided. However, once the required inputs were prepared, the tool substantially reduced the time and manual effort needed to generate submission-ready *Assessments* and *Lab Tests* templates while producing outputs that were consistent with ImmPort curation practices and repository formatting requirements.

## Discussion

The widespread push for open science and data transparency has underscored the importance of not only sharing data but also ensuring that shared datasets are well annotated and reusable^38–40^. For mechanistic and clinically focused immunological research, repositories like ImmPort play a pivotal role in enabling secondary analyses, meta-studies, and reproducibility. However, despite ImmPort providing structured templates and guides to support standardized data submission, their consistent use remains a challenge. This is particularly evident in the underutilization of the *Assessments* and *Lab Tests* templates, which are essential for capturing clinical and phenotypic data and ensuring data completeness.

The ImmPort Data Curation Tool is designed to address this gap by simplifying the process of *Assessments* and *Lab Tests* template generation. Through a semi-automated workflow and a user-friendly GUI, the tool reduces the technical expertise and manual effort required by data contributors. It has been noted that easy-to-use tools that partially automate data curation help researchers create and share structured data, and this curation tool reflects this approach^39^. Moreover, such automation can also benefit the repository curators — either directly, by supporting their workflows, or indirectly, by improving the quality and consistency of submitted data and thereby reducing the effort required for post-submission curation^41^.

The tool was evaluated across seven different ImmPort studies, resulting in completed *Assessments* and *Lab Tests* templates that averaged thousands of rows and columns. The templates exhibited substantial data volume and notable interstudy variability, reflecting both the complexity of the template formats and the study-specific differences, including study size and design. Ultimately, the completed templates demonstrated that the tool can reliably produce submission-ready templates that conform to ImmPort’s formatting standards and pass internal validation. This not only improves data completeness but also enhances the overall quality and interoperability of the repository. This is especially important for studies that include IPD, where detailed and standardized data are essential for enabling robust secondary analyses^11^.

A key feature of the tool is its accessibility to users without programming expertise. By leveraging a GUI, the tool allows a broad range of researchers without programming experience to contribute high-quality data. At present, the requirement for a curated data dictionary introduces a manual step that can be time-consuming and is potentially dependent on the curator’s familiarity with the study. While many clinical studies already generate data dictionaries, those data dictionaries are often provided in inconsistent file formats and may be poorly formatted or missing important data^42–44^. Even once issues are fixed and the dictionary is aligned withthe tool’s required format, the added columns require additional effort. While most additions to the data dictionary are still user-optional, compliance is encouraged because it ultimately improves the completeness of the templates. This step currently represents the primary bottleneck in the workflow. However, it should be noted that the seven studies tested here were done by individuals that were not part of the original study teams and whose knowledge of each study was obtained from documents and files within ImmPort. This illustrates the feasibility of using this tool even with data from studies that users have limited prior familiarity.

One of the most impactful future enhancements to the ImmPort Data Curation Tool would be reducing or eliminating the manual effort required to curate the data dictionary. Techniques from natural language processing, including large language models, could be employed to extract information from CRFs, study protocols, and other study documentation and incorporate that information into the data dictionary. Such approaches would require validation and quality control procedures to ensure accuracy and formatting consistency. This would significantly reduce the manual workload and further streamline the curation process. Additionally, integrating structured, semantic representations to the tool-specific curated fields in the data dictionary could enhance the accuracy and consistency of the dictionary itself, improve the quality of the generated templates, and ultimately strengthen the standardized data within ImmPort^44^.

Another limitation of the current tool is its scope. The focus on *Assessments* and *Lab Tests* is justified by their low completion rates, high informational value, and applicability to a large percentage of study files (51.3% of uploaded study files from the seven evaluated ImmPort studies). However, ImmPort also includes several other templates that are important for comprehensive data annotation. These include templates for demographics, interventions, concomitant medications, adverse events, and assay-specific data (e.g., ELISA, flow cytometry, and gene expression templates). Expanding the tool to support these additional templates would provide a more inclusive solution and further promote adherence to FAIR data principles.

A key strength of this semi-automated curation approach is its potential applicability to other data repositories. Although this tool was developed specifically for ImmPort, the underlying approach — combining a curated data dictionary with study-specific data files for a semi-automated template generation process — could be adapted for other repositories that encourage or require structured data submissions. This makes it a broadly applicable solution for improving structured data quality across platforms.

In conclusion, the ImmPort Data Curation Tool represents a significant step toward improving data quality in immunological research. By reducing the barriers to template completion, it encourages broader participation in data sharing and enhances the utility of public datasets. Continued development and expansion of the tool’s capabilities will further support the goals of open science and data-driven discovery.

## Data Availability

The datasets analyzed during this study are publicly available in the ImmPort repository (https://immport.org) under the following accession numbers: SDY620, SDY1027, SDY1433, SDY1515, SDY1519, SDY1520, and SDY1550.

## Code Availability

The ImmPort Data Curation Tool is openly available on GitHub (https://github.com/AndorfLab/ImmPort-Curation-Tool), along with a full list of dependencies and installation instructions.

## Author Contributions

A.R.T., J.D.F., and K.B. developed the methodology and software and performed data curation. A.R.T. and J.D.F. prepared the online user guide. A.R.T. conducted data analysis and prepared the original manuscript draft. N.K.S. tested the tool and aided in data curation. P.P., M.D.S., P.D., and H.S. provided expertise in ImmPort data standards and workflows. P.P. and M.D.S evaluated the tool, contributed to the Usability Evaluation section of the manuscript, and uploaded the completed templates to ImmPort. P.D. and H.S. supported the development of the methodology. S.A. conceived and supervised the study, interpreted results, and edited the manuscript. All authors reviewed and approved the final manuscript.

## Acknowledgements

We thank the ImmPort team for their valuable discussions, feedback, and data upload assistance throughout this study. We also thank those that tested the application and provided valuable feedback.

