## Supplement for "A semi-automated tool for improving *Assessments* and *Lab Tests* data annotation in ImmPort"

**Methods**

**Assignments in the “Mappings” column of the data dictionary**

The "Mappings" column can be filled in with one of six options – “User Defined ID”, “Study Day”, “Study Time”, “Visit”, “Category”, or “NA” – or left empty entirely. If it is left empty, that “Field Name” will be its own entry in the completed template. In other words, the values from the corresponding column in the study file will be used to propagate the “Result Value Reported” field. It’s expected that “Mappings” will be blank for most fields, since the majority of columns in a study file will contain results.

Alternatively, “User Defined ID” should be used to indicate the "Field Name" that contains the participant ID. “Visit” should be used to indicate the "Field Name" that contains the specific visit information. “Study Day” should be used to indicate the day, while “Study Time” should be used to indicate the time. All these designations should only be used a maximum of once per "Table Name". If there are multiple study days or times represented within a table, the "Override Study Day" and "Override Study Time" columns in the data dictionary should also be used.

The other “Mappings” options, “NA” and “Category”, can be used more than once per "Table Name". “NA” should be used to indicate that the "Field Name" should be ignored entirely from the completed *Lab Tests* or *Assessments* template. For example, if a "Field Name" is only used for internal record keeping and is meaningless to external researchers, then that "Field Name" should be designated as “NA”.

“Category” should be used when that "Field Name" does not contain a result per se, but rather a category that adds specificity to the results. That value will then be added to all of the "Field Descriptions" for that "Table Code" instead of being listed as a result in the final template. For example, consider a "Field Name" that is SKPT with a "Field Description" of SKIN PRICK TEST RESULT and with values in the study file containing those measurements (e.g., 1, 3, 0, 4). Consider another "Field Name" for that table is ALLER with a "Field Description" of ALLERGENS, containing different types of allergens in the study file (e.g., peanut, pollen, milk). The ALLER field might be better designated as “Category”. That would then make SKPT’s "Field Description" correspond to the data. In other words, it might now be PEANUT SKIN PRICK TEST RESULT, POLLEN SKIN PRICK TEST RESULT, MILK SKIN PRICK TEST RESULT, etc.

“User Defined ID”, “Study Day”, “Visit”, “Category”, and “NA” are all used to propagate both the *Lab Tests* and *Assessments* templates. “Study Time” is only used within the *Assessments* template.

**Tables**

Table S1. Mapping of each column in the generated *Assessments* template to its corresponding source from the uploaded ImmPort-generated files, the curated data dictionary, inputs within the GUI, the study file itself, or unique values generated by the ImmPort Data Curation Tool. The ‘Explanation’ column summarizes how the template column is completed. When an “OR” is included, the top-to-bottom order reflects the priority in which each source is used (i.e., if a value is present in “Override Study Time,” it will be used before a ‘Study Time’ value in “Mappings” is used).

| **Template**  **Column** | **Source of Value** | | | | **Explanation** |
| --- | --- | --- | --- | --- | --- |
|  | **ImmPort-generated Files** | **Data Dictionary** | **GUI**  **Input** | **Study Files** |  |
| Subject ID |  | X |  | X | “Mappings” column (specified as ‘User Defined ID’) in the data dictionary, and study file |
| Assessment Panel ID |  |  |  |  | Unique value generated by the tool |
| Study ID | X |  |  |  | "STUDY_ACCESSION" column from the uploaded ImmPort-generated study data file (study_data.txt) |
| Name Reported | X |  |  |  | "DESCRIPTION" column from the uploaded ImmPort-generated study data file (study_data.txt) |
| Assessment Type |  |  | X |  | User specified within the curation tool ("Assessment Name" textbox) |
| Status |  |  |  |  | NA* |
| CRF File Names |  |  |  | X | Study file name |
| User Defined ID |  |  |  |  | Unique value generated by the tool |
| Planned Visit ID | X | X | X | X | “Mappings” column (specified as ‘Visit’) in the data dictionary, uploaded ImmPort-generated planned visit file (planned_visit.txt), and study file  OR  User specified within the curation tool ("Default Visit" dropdown) and uploaded ImmPort-generated planned visit file (planned_visit.txt) |
| Name Reported |  | X |  |  | "Field Description" column in the data dictionary |
| Study Day | X | X |  | X | "Override Study Day" column in the data dictionary, and study file  OR  "Override Study Day" column in the data dictionary**  OR  “Mappings” column (specified as ‘Study Day’) in the data dictionary, and study file  OR  “MIN_START_DAY” column from the uploaded ImmPort-generated planned visit file (planned_visit.txt) that corresponds to the specified “visit” |
| Age at Onset Reported |  | X |  | X | "Age at Onset Reported" column in the data dictionary, and study file  OR  "Age at Onset Reported" column in the data dictionary** |
| Age at Onset Unit Reported |  | X |  | X | "Age at Onset Unit Reported" column in the data dictionary, and study file  OR  "Age at Onset Unit Reported" column in the data dictionary** |
| Is Clinically Significant |  |  |  |  | NA* |
| Location of Finding Reported |  |  |  |  | NA* |
| Organ or Body System Reported |  | X |  | X | "Location" column in the data dictionary, and study file  OR  "Location" column in the data dictionary** |
| Result Value Reported |  | X |  | X | Columns in the study file not specified in the “Mappings” column as ‘User Defined ID’, ‘Study Day’, ‘Study Time’, ‘Visit’, ‘Category’, or ‘NA’ |
| Result Unit Reported |  | X |  | X | "Unit" column in the data dictionary, and study file  OR  "Unit" column in the data dictionary** |
| Result Value Category |  |  |  |  | NA* |
| Subject Position Reported |  |  |  |  | NA* |
| Time of Day |  | X |  | X | "Override Study Time" column in the data dictionary, and study file  OR  "Override Study Time" column in the data dictionary**  OR  “Mappings” column (specified as ‘Study Time’) in the data dictionary, and study file |
| Verbatim Question |  | X |  | X | "Verbatim Question" column in the data dictionary, and study file  OR  "Verbatim Question" column in the data dictionary** |
| Who is Assessed |  | X |  | X | "Who is Assessed" column in the data dictionary, and study file  OR  "Who is Assessed" column in the data dictionary** |

* NA values denote template columns that are not propagated by the ImmPort Curation Tool.

** The curation tool first checks whether the value specified in the data dictionary corresponds to another “Field Name” in the data dictionary. If it does, the values from that field in the study file are used. Otherwise, the literal value provided in the data dictionary is applied.

Table S2. Mapping of each column in the generated *Lab Tests* template to its corresponding source from the uploaded ImmPort-generated files, the curated data dictionary, inputs within the GUI, the study file itself, or unique values generated by the ImmPort Data Curation Tool. The ‘Explanation’ column summarizes how the template column is completed. When an “OR” is included, the top-to-bottom order reflects the priority in which each source is used (i.e., if a ‘Visit’ value is present in “Mappings”, it will be used before a "Default Visit" that is specified within the curation tool is used).

| **Template Column** | **Source of Value** | | | | **Explanation** |
| --- | --- | --- | --- | --- | --- |
|  | **ImmPort-generated Files** | **Data Dictionary** | **GUI**  **Input** | **Study Files** |  |
| Biosample ID |  |  |  |  | Unique value generated by the tool |
| Lab Test Panel ID |  |  |  |  | Unique value generated by the tool |
| Study ID | X |  |  |  | "STUDY_ACCESSION" column from the uploaded ImmPort-generated study data file (study_data.txt) |
| Protocol ID(s) | X |  | X |  | User specified within the curation tool (“Protocol” dropdown), and "PROTOCOL_ACCESSION" column from the uploaded ImmPort-generated protocol file (protocol.txt) |
| Subject ID |  | X |  | X | “Mappings” column (specified as ‘User Defined ID’) in the data dictionary, and study file |
| Planned Visit ID | X | X | X | X | “Mappings” column (specified as ‘Visit’) in the data dictionary, uploaded ImmPort-generated planned visit file (planned_visit.txt), and study file  OR  User specified within the curation tool ("Default Visit" dropdown), and uploaded ImmPort-generated planned visit file (planned_visit.txt) |
| Type |  |  | X |  | User specified within the curation tool (“Type” dropdown) |
| Subtype |  |  | X |  | User specified when "Type" = "Other" within the curation tool ("Subtype" textbox) |
| Name |  |  |  |  | NA* |
| Description |  |  |  |  | NA* |
| Study Time Collected | X | X |  | X | "Override Study Day" column in the data dictionary, and study file  OR  "Override Study Day" column in the data dictionary**  OR  “Mappings” column (specified as ‘Study Day’) in the data dictionary, and study file  OR  “MIN_START_DAY” column from the uploaded ImmPort-generated planned visit file (planned_visit.txt) that corresponds to the specified “visit” |
| Study Time Collected Unit |  |  |  |  | “Days” |
| Study Time T0 Event |  |  | X |  | User specified within the curation tool ("Study Time T0 Event" dropdown) |
| Study Time T0 Event Specify |  |  | X |  | User specified when "Study Time T0 Event" = "Other" within the curation tool ("Study Time T0 Event Specify" textbox) |
| Name Reported |  |  | X |  | User specified within the curation tool ("Name Reported" dropdown) |
| User Defined ID |  |  |  |  | Unique value generated by tool |
| Name Reported*** |  | X |  |  | "Field Description" column in the data dictionary |
| Result Value Reported |  | X |  | X | Columns in the study file not specified in the “Mappings” column as ‘User Defined ID’, ‘Study Day’, ‘Study Time’, ‘Visit’, ‘Category’, or ‘NA’ |
| Result Unit Reported |  | X |  | X | "Unit" column in the data dictionary, and study file  OR  "Unit" column in the data dictionary** |

* NA values denote template columns that are not propagated by the ImmPort Curation Tool.

** The curation tool first checks whether the value specified in the data dictionary corresponds to another “Field Name” in the data dictionary. If it does, the values from that field in the study file are used. Otherwise, the literal value provided in the data dictionary is applied.

*** There are two “Name Reported” columns in the *Lab Tests* template. The first “Name Reported” column should describe the specific lab test, while the second “Name Reported” column should describe the “Result Value Reported”.

Table S3. Study files that were excluded from transformation into the *Assessments* or *Lab Tests* templates. These files were omitted either because they contained administrative data or were better suited to other ImmPort templates (i.e., demographics, interventions, concomitant medications, adverse events, and assay-specific templates).

| **ImmPort Study**  **Accession** | **Excluded Study Files** | **Number of Excluded Study Files** |
| --- | --- | --- |
| SDY620 | AEBL – Adverse events – baseline visit  AE28 – Adverse events – day 28  AE37 – Adverse events – day 3 and day 7  AETA – Adverse events – telephone assessment  AEXP – Adverse events  BLBR – Blinding  CMED – Concomitant medications log  CMYN – Concomitant Medications Yes/No  DEMO – Demographics  ELIG – Eligibility checklist  RECR – Recruitment  RESC – Rescheduling  SAEV – Serious adverse event  SUBJ – Participant ID  VACC – Vaccine administration | 15 |
| SDY1027 | ACI – Additional contact information ACIU – Additional contact information update AE – Adverse events review AEA – Adverse events assessment BSCFE – Family blood draw willingness CM – Concomitant and prescribed medications review CMA – Concomitant medications assessment CN – Administrative DF – Demographics DPR – Discontinuation prior to screening ISI – ICAC2 study ID RF – Recruitment RL – Reimbursement SEC - Eligibility Checklist SEC2 - Eligibility Checklist Continuation | 15 |
| SDY1433 | ADDP – Serious Adverse Event AEXP – Adverse Event AUTOAB – AutoAB BGE – mRNA Blood CM01 - Concomitant medications CM02 - Concomitant medications CM03 - Concomitant medications CM04 - Concomitant medications CM05 - Concomitant medications CM06 - Concomitant medications CM07 - Concomitant medications CM08 - Concomitant medications CM09 - Concomitant medications CM10 - Concomitant medications CM12 - Concomitant medications CM14 - Concomitant medications CM15 - Concomitant medications DDEM – Donor demographics DEMO – Recipient demographics DETH – Death ELIG - Eligibility GENR – DNA Genomics-Recipient HLAAB – Anti-HLA AB HOSP – Hospitalization IMDM – Infection prophylaxis and treatment IMED – Maintence immunosuppressive medications INFC – Infection IPMD – Maintence immunosuppressive medications ISIT – Infection site METH – Medrol (methylprednisolone) or equivalent log NUL2 – Nulojix (belatacept) vitals NULO – Nulojix (belatacept) vitals PGRF – Pancreatic graft failure REIN – Reinstatement of tacrolimus SAEV – Serious Adverse Event TACW – Initiation/completion of tacrolimus withdrawal TGE – mRNA Tissue THYM – Thymoglobulin log USP – ELISA | 39 |
| SDY1515 | AD1 – Adverse event AD2 – Adverse event AD3 – Adverse event CMD – Medication DEM – Demographics DS4 – Dosing symptom log ECCF4A – Eligibility ECCF4B – Eligibility ECCF4M – Eligibility ENROLL – Enrollment date FAE – Food allergy event PDV – Non subject specific protocol deviation PT4 – Study product tracking – CoFAR4 | 13 |
| SDY1519 | DEM – Demographics ECCFA – Consent for future specimen storage, check off inclusion/exclusion criteria ENROLL – Enrollment date, protocol segment PDV – Non subject specific protocol deviation | 4 |
| SDY1520 | AD1 – Adverse event  AD2 – Adverse event  AD3 – Adverse event  DEM – Demographics  DS6 – Dosing symptom log  DSL_MEDDRA – Dosing  ECCF6A – Informed consent, data storage  ECCF6B – Eligibility  ENROLL – Enrollment date  FAE – Food allergy episode  MED – Medication  PDV – Non subject specific protocol deviation  PT6 – Study product tracking - CoFAR6  TRTGRP – Treatment group  WHOCF6 – WHO medication | 15 |
| SDY1550 | AD1 – Adverse event  AD2 – Adverse event  AD3 – Adverse event  DEM – Demographics  DSL – Dosing symptom log  DSL_MEDDRA – Dosing symptom MedDRA  ECCF7A – Informed consent, data storage  ECCF7B – Eligibility  ENROLL – Enrollment date  ESC – Initial Escalation  FAE – Food allergy episode  MED – Medication  PDV – Non subject specific protocol deviation  PTB – Study product tracking (build-up)  PTC – Study product tracking (bulk powder)  WHOCF7 – WHO medication | 16 |
| **Total** | | 117 |

Table S4. Study files from ImmPort study SDY620 and the resulting number of rows and columns in the generated templates. The total number of rows and columns are shown (including empty cells). Additionally, the median number of columns filled per row and the first (Q1) and third (Q3) quartiles are reported.

| ***Assessments* Template** | | | |
| --- | --- | --- | --- |
| **Case Report Form** | **Number of**  **Rows** | **Number of Columns** | **Median Number of Columns**  **(Q1, Q3)** |
| **COMP – Study Completion Final Assessment** | 361 | 169 | 69 (61, 69) |
| **DEVI – Protocol Deviation** | 77 | 233 | 55 (48, 55) |
| **DISC – Study Discontinuation** | 16 | 41 | 20 (20, 20) |
| **EASI – EASI Score** | 233 | 169 | 46 (46, 46) |
| **MDHX – Medical History** | 143 | 105 | 48 (48, 48) |
| **PHEX – Physical Exam** | 372 | 201 | 53 (29, 53) |
| **RLSC – Rajika-Langeland Score** | 235 | 137 | 38 (38, 38) |
| **VITL – Vital Signs** | 371 | 201 | 53 (53, 53) |
| ***Lab Tests* Template** | | | |
| **BS28 – Blood Sample Collection – Day 28/Unscheduled Visit** | 361 | 45 | 30 (30, 33) |
| **BSBL – Blood Sample Collection – Baseline Visit** | 421 | 65 | 42 (42, 48) |
| **BSCR – Blood Sample Collection – Cellular Immune Response** | 204 | 41 | 30 (30, 30) |
| **BSMC – Blood Sample Collection – Microarray** | 185 | 57 | 39 (33, 42) |
| **CBCV – CBC Sample Results** | 247 | 61 | 15 (15, 15) |
| **PREG – Pregnancy Test Results** | 217 | 29 | 21 (21, 21) |

Table S5. Study files from ImmPort study SDY1027 and the resulting number of rows and columns in the generated templates. The total number of rows and columns are shown (including empty cells). Additionally, the median number of columns filled per row and the first (Q1) and third (Q3) quartiles are reported.

| ***Assessments* Template** | | | |
| --- | --- | --- | --- |
| **Case Report Form** | **Number of**  **Rows** | **Number of Columns** | **Median Number of Columns**  **(Q1, Q3)** |
| **ASTR - Allergen Skin Test Results** | 200 | 1,929 | 602 (244.5, 642) |
| **HE - Home Environment** | 200 | 937 | 212 (204, 228) |
| **MC - Methacholine Challenge** | 34 | 761 | 395 (375, 415) |
| **MCOR – Methacholine Challenge** | 34 | 313 | 58 (51, 58) |
| **PDR – Protocol Deviation Report** | 12 | 201 | 53 (53, 53) |
| **PE - Vital Signs, Growth Parameters and Physical Exam** | 200 | 3,529 | 448 (442, 458) |
| **PEU - Vital Signs, Growth Parameters and Physical Exam Update** | 36 | 521 | 149 (149, 149) |
| **PX - Participant Exposures** | 200 | 745 | 134 (125, 188) |
| **RHQ - Respiratory Health Questionnaire – Participant** | 99 | 1,481 | 473 (458, 493) |
| **SCS – Study Completion Status** | 200 | 73 | 21 (21, 21) |
| **SOR – Spirometry** | 200 | 393 | 59 (59, 96) |
| **ST – Spirometry** | 200 | 745 | 189 (169, 279) |
| ***Lab Tests* Template** | | | |
| **BSCFM - Blood Sample Collection – Family Member** | 56 | 129 | 114 (110, 114) |
| **BSCP - Blood Sample Collection – Participant** | 204 | 61 | 45 (42, 45) |
| **IGEL – IgE** | 200 | 53 | 37 (37, 41) |
| **NECSC - Nasal Epithelial Cell Sample Collection** | 102 | 41 | 36 (36, 36) |
| **PTR - Pregnancy Test Results** | 146 | 29 | 16 (16, 16) |

Table S6. Study files from ImmPort study SDY1433 and the resulting number of rows and columns in the generated templates. The total number of rows and columns are shown (including empty cells). Additionally, the median number of columns filled per row and the first (Q1) and third (Q3) quartiles are reported.

| ***Assessments* Template** | | | |
| --- | --- | --- | --- |
| **Case Report Form** | **Number of Rows** | **Number of Columns** | **Median Number of Columns**  **(Q1, Q3)** |
| **DEVI – Protocol Deviation** | 44 | 3,145 | 407 (288.75, 574.5) |
| **DISP – Final Disposition** | 44 | 57 | 20.5 (13, 21) |
| **MDHX – Medical History** | 43 | 425 | 133 (133, 180.5) |
| **MOTH – Medical History** | 31 | 809 | 76 (48, 107.5) |
| **NEUR – Neurological Exam for PML** | 43 | 3,721 | 1,173 (1,166, 1,405) |
| **PBIO – Local Pancreatic Biopsy/Suspected Rejection** | 19 | 249 | 54 (33.5, 90) |
| **RBIO – Local Renal Biopsy/Suspected Rejection** | 21 | 489 | 70 (54, 102) |
| **TDIS – Study Therapy Discontinuation, Reduced Follow-up** | 14 | 169 | 74 (74, 74) |
| **TRAN – Transplant Donor** | 43 | 1,081 | 298 (282, 362) |
| **TRN2 – Transplant Recipient** | 43 | 1,369 | 477 (442.5, 536.5) |
| **VIS2 – Study Visit Form 1** | 43 | 4,809 | 1,558 (1,419, 1,949.5) |
| **VIS3 – Study Visit Form 2** | 43 | 249 | 111 (111, 111) |
| **VITL - Physical Exam** | 43 | 137 | 74 (74, 74) |
| ***Lab Tests* Template** | | | |
| **CHEM - Serum Chemistry** | 402 | 77 | 68 (47, 69) |
| **CPEP - C-Peptide** | 76 | 25 | 15 (15, 15) |
| **FAST - Fasting Lipid Panel** | 154 | 37 | 30 (30, 30) |
| **HEMA - Hematology** | 329 | 77 | 66 (66, 67) |
| **HEMO - Hemoglobin A1c** | 280 | 21 | 15 (15, 15) |
| **IMDL - Immunosuppressive Medication Trough Levels** | 734 | 37 | 27 (27, 27) |
| **PRG2 – Pregnancy Test** | 43 | 25 | 14 (14, 14) |
| **TRN4 - Recipient CMV, BKV by PCR** | 199 | 37 | 17 (17, 25) |
| **URIN - Urinalysis** | 323 | 33 | 23 (19, 23) |

Table S7. Study files from ImmPort study SDY1515 and the resulting number of rows and columns in the generated templates. The total number of rows and columns are shown (including empty cells). Additionally, the median number of columns filled per row and the first (Q1) and third (Q3) quartiles are reported.

| ***Assessments* Template** | | | |
| --- | --- | --- | --- |
| **Case Report Form** | **Number of Rows** | **Number of Columns** | **Median Number of Columns**  **(Q1, Q3)** |
| **ADF – Atopic Dermatitis** | 50 | 2,681 | 384 (201.75, 574.5) |
| **DEV – Protocol Deviation** | 27 | 1,161 | 153 (83, 233.5) |
| **DM2 – Protocol Demographics** | 62 | 89 | 40 (40, 40) |
| **EET – Endpoint Titration** | 62 | 1,353 | 382 (194, 574.5) |
| **ES4 – Escalation** | 40 | 8,313 | 2,344.5 (1,575.25, 2,883.75) |
| **MD1 – Baseline Medical History Part I** | 61 | 777 | 301 (224, 358) |
| **MD4 – Baseline Medical History Part II** | 61 | 1,593 | 654 (572, 751) |
| **MD3 – Baseline Medical History Part III** | 61 | 601 | 249 (233, 267) |
| **OFC – Oral Food Challenge** | 48 | 3,705 | 995 (484.5, 1,282.75) |
| **PDD – Permanent Dosing Discontinuation** | 40 | 57 | 26 (20, 26) |
| **SPT – Prick Skin Results** | 62 | 505 | 219 (157, 219) |
| **VS3 – Visit Form Part I** | 55 | 18,873 | 4,354 (249.5, 6,275.5) |
| **VS4 – Visit Form Part II** | 56 | 2,361 | 230 (69, 312.25) |
| **WDF – Withdrawal/Termination Form** | 62 | 89 | 29 (22, 37) |
| ***Lab Tests* Template** | | | |
| **BAS – Basophil** | 52 | 4,793 | 2,985 (1,391, 3,678) |
| **IGE – Antibody Assay Results** | 384 | 97 | 25 (25, 39) |
| **LB4 – CoFAR4 Lab Form** | 326 | 405 | 312 (73, 359) |
| **PRO – Proliferation of 3H-Thymidine** | 203 | 177 | 131 (71, 131) |

Table S8. Study files from ImmPort study SDY1519 and the resulting number of rows and columns in the generated templates. The total number of rows and columns are shown (including empty cells). Additionally, the median number of columns filled per row and the first (Q1) and third (Q3) quartiles are reported.

| ***Assessments* Template** | | | |
| --- | --- | --- | --- |
| **Case Report Form** | **Number of Rows** | **Number of Columns** | **Median Number of Columns**  **(Q1, Q3)** |
| **DEV – Protocol Deviation** | 93 | 345 | 73 (73, 79) |
| **DM2 – Protocol Demographics** | 708 | 153 | 77 (77, 77) |
| **EDO – Endoscopy Form** | 562 | 153 | 69 (68, 69) |
| **EE1 – EE Baseline Medical History** | 708 | 633 | 325 (321, 330) |
| **EE2 – Symptoms** | 708 | 457 | 195 (184.75, 213) |
| **EE3 – Allergies Part I** | 708 | 873 | 112 (24, 155.25) |
| **EE4 – Allergies Part II** | 707 | 745 | 85 (42, 112) |
| **EE5 – Surgical History** | 708 | 169 | 69 (69, 69) |
| **EE6 – Family History Part I** | 708 | 601 | 79.5 (57, 111) |
| **EE7 – Family History Part II** | 680 | 137 | 30 (30, 30) |
| **WDF – Withdrawal/Termination Form** | 5 | 73 | 29 (29, 29) |
| ***Lab Tests* Template** | | | |
| **EEP – Pathology Form** | 708 | 113 | 44 (36, 59) |
| **LB5 – CoFAR5 Lab Form** | 1,038 | 45 | 19 (18, 25) |

Table S9. Study files from ImmPort study SDY1520 and the resulting number of rows and columns in the generated templates. The total number of rows and columns are shown (including empty cells). Additionally, the median number of columns filled per row and the first (Q1) and third (Q3) quartiles are reported.

| ***Assessments* Template** | | | |
| --- | --- | --- | --- |
| **Case Report Form** | **Number of Rows** | **Number of Columns** | **Median Number of Columns**  **(Q1, Q3)** |
| **ADF – Atopic Dermatitis** | 86 | 4,393 | 493.5 (357, 1,095.25) |
| **DEV – Protocol Deviation** | 69 | 2,025 | 241 (89, 386) |
| **DM2 – Protocol Demographics** | 86 | 121 | 59 (59, 59) |
| **MD1 – Baseline Medical History Part I** | 86 | 825 | 311 (260.25, 363.25) |
| **MD4 – Baseline Medical History Part II** | 86 | 1,657 | 624 (546.75, 691.25) |
| **MD3 – Baseline Medical History Part III** | 86 | 601 | 249 (233, 267) |
| **OFC – Oral Food Challenge** | 84 | 3,881 | 1,299 (887.5, 1,394) |
| **PDD – Permanent Dosing Discontinuation** | 75 | 73 | 29 (29, 29) |
| **PEX – Physical Examination** | 86 | 873 | 361 (221.75, 386) |
| **SAF – Skin Assessment Form** | 71 | 1,737 | 134 (86, 210.5) |
| **SPT – Prick Skin Results** | 86 | 2,985 | 1,160 (871.5, 1,240) |
| **VS3 – Visit Form Part I** | 85 | 10,233 | 3,202 (2,591, 4,003) |
| **VS4 – Visit Form Part II** | 71 | 4,825 | 649 (302.5, 1,232) |
| **WDF – Withdrawal/Termination Form** | 86 | 105 | 22 (22, 29) |
| ***Lab Tests* Template** | | | |
| **BAS – Basophil** | 383 | 1,505 | 507 (487, 515) |
| **FCY – Flow Cytometry** | 236 | 4,733 | 2,195 (2,079, 4,154) |
| **IGE – Antibody Assay Results** | 1,547 | 57 | 32 (25, 32) |
| **LB6 – CoFAR6 Lab Form** | 548 | 113 | 58 (55, 74) |
| **UPG – Urinalysis/Pregnancy/Guaiac** | 80 | 25 | 15 (15, 15) |

Table S10. Study files from ImmPort study SDY1550 and the resulting number of rows and columns in the generated templates. The total number of rows and columns are shown (including empty cells). Additionally, the median number of columns filled per row and the first (Q1) and third (Q3) quartiles are reported.

| ***Assessments* Template** | | | |
| --- | --- | --- | --- |
| **Case Report Form** | **Number of Rows** | **Number of Columns** | **Median Number of Columns**  **(Q1, Q3)** |
| **ADF – Atopic Dermatitis** | 187 | 1,945 | 109 (33, 285) |
| **DEV – Protocol Deviation** | 69 | 2,649 | 174 (90, 328) |
| **DM2 – Protocol Demographics** | 187 | 121 | 52 (52, 52) |
| **ENC – Study Encounter** | 113 | 9,481 | 1,706 (192, 3,405) |
| **LF7 – Long Term Follow-up** | 67 | 2,329 | 477 (318, 565) |
| **MD1 – Baseline Medical History Part I** | 187 | 1,417 | 298 (262, 349) |
| **MD4 – Baseline Medical History Part II** | 187 | 2,105 | 583 (500.5, 681.5) |
| **MD3 – Baseline Medical History Part III** | 187 | 1,065 | 240 (233, 267) |
| **OFC – Oral Food Challenge** | 169 | 3,833 | 578 (428, 1,312) |
| **PDD – Permanent Dosing Discontinuation** | 92 | 57 | 21 (21, 21) |
| **PEX – Physical Examination** | 187 | 1,673 | 212 (188, 604) |
| **SPT – Prick Skin Results** | 187 | 1,689 | 255 (255, 629) |
| **VS3 – Visit Form Part I** | 90 | 1,961 | 810.5 (655.25, 918.5) |
| **VS4 – Visit Form Part II** | 91 | 2,057 | 262 (182, 469) |
| **WDF – Withdrawal/Termination Form** | 187 | 105 | 37 (22, 37) |
| ***Lab Tests* Template** | | | |
| **BAS – Basophil** | 145 | 4,437 | 2,395 (487, 3,735) |
| **FCY – Flow Cytometry** | 143 | 11,989 | 6,731 (1,691, 8,994.5) |
| **IGE – Antibody Assay Results** | 1,936 | 97 | 25 (18, 32) |
| **LB7 – CoFAR7 Lab Form** | 665 | 89 | 63 (63, 66) |
| **UPG – Urinalysis/Pregnancy/Guaiac** | 20 | 25 | 15 (15, 15.75) |

**Figures**

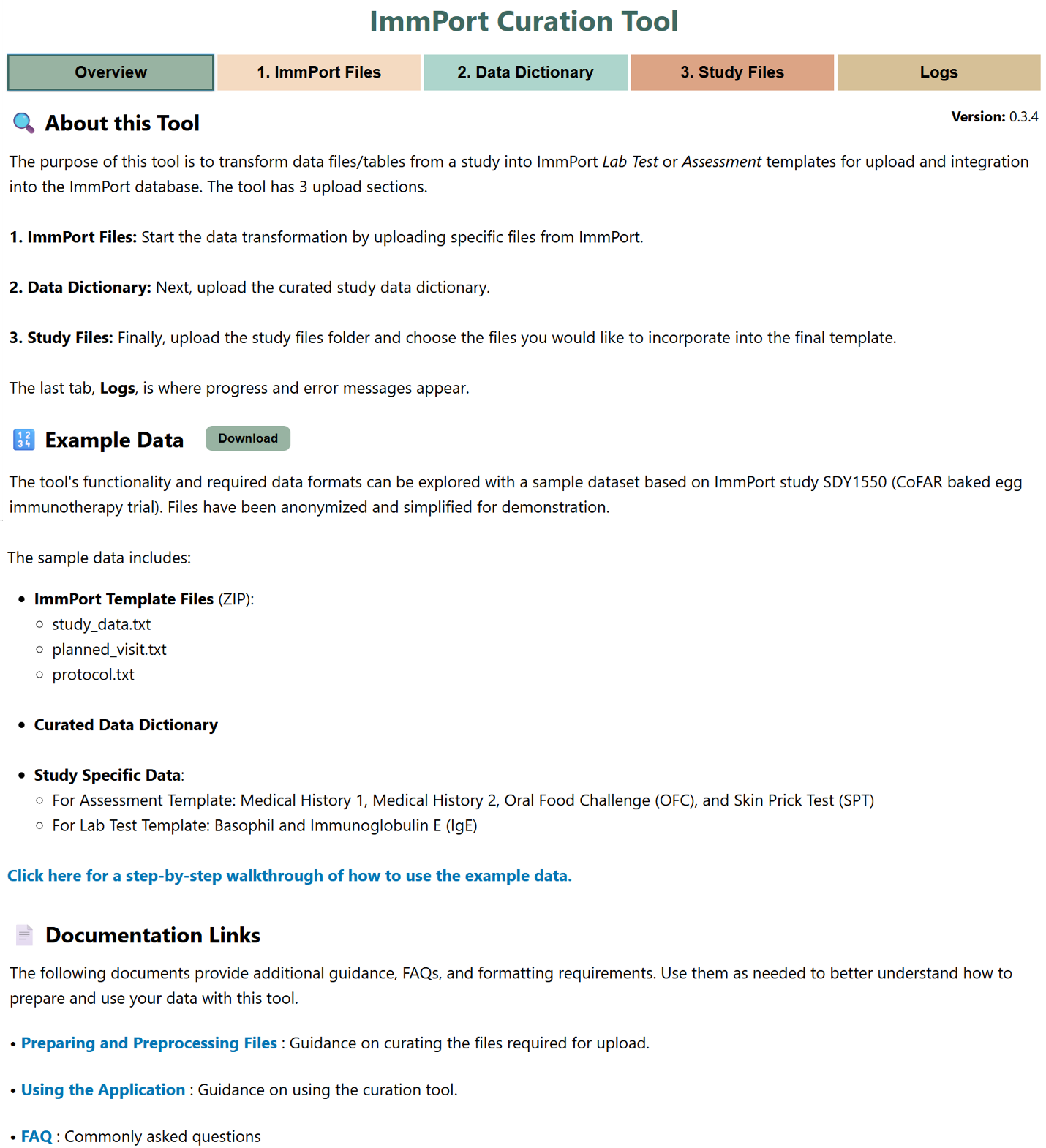

Figure S1. Screenshot of the “Overview” tab in the GUI of the curation tool. This tab provides a summary of the tool’s functionality, a direct download link to example data, and access to extended documentation. It serves as a starting point for users to understand the workflow and available resources.

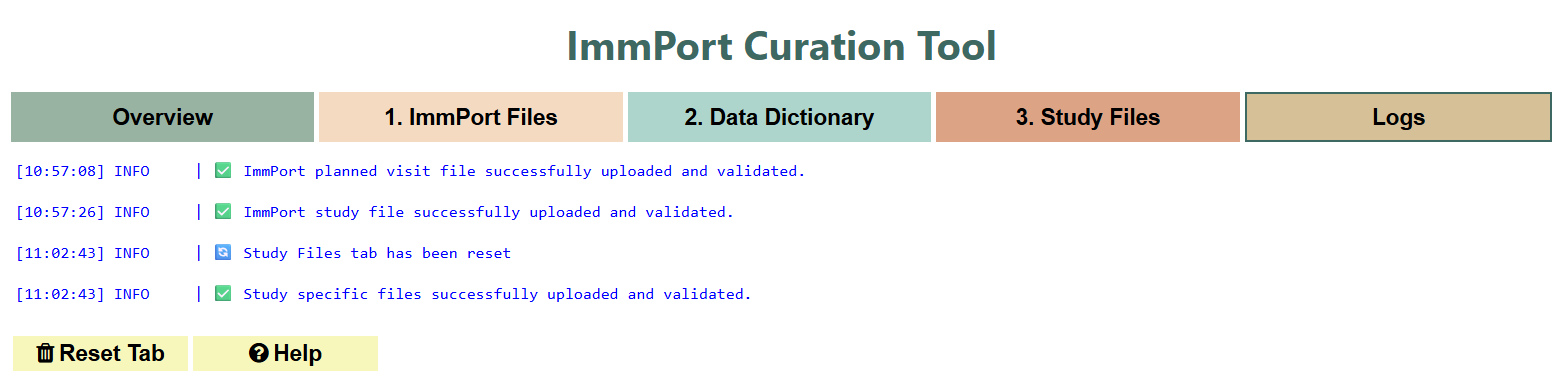

Figure S2. Screenshot of the “Logs” tab in the GUI of the curation tool. This tab displays real-time status updates and error messages, helping users monitor progress and troubleshoot issues during file uploads and template generation. Example logs are shown in the application for demonstration purposes.
